# Time-resolved, single-molecule, correlated chemical probing of RNA

**DOI:** 10.1101/2020.03.21.001560

**Authors:** Jeffrey E. Ehrhardt, Kevin M. Weeks

## Abstract

Methods for capturing the folding dynamics of functionally important RNAs, especially large RNAs, have relied primarily on global measurements of structure or on per-nucleotide chemical probing. These approaches infer, but do not directly measure, through-space tertiary interactions. Here we introduce trimethyloxonium (TMO) as a chemical probe for RNA. TMO enables time-resolved, single-molecule, through-space structure probing of RNA folding using a correlated chemical probing framework. TMO methylates RNA about 90 times faster than the widely used dimethyl sulfate probe, allowing structure interrogation on the second time scale. We used TMO to monitor folding of the RNase P RNA – a functional RNA with extensive long-range and noncanonical interactions – by direct measurement of through-space tertiary interactions in a time-resolved way. Time-dependent correlation changes directly revealed the central role of a long-range tertiary loop-loop interaction that guides native RNA folding. Single-molecule, time-resolved RNA structure probing with TMO is poised to reveal a wide range of dynamic RNA folding processes and principles of RNA folding.

## Introduction

Chemical probing strategies are widely and effectively used to monitor RNA folding reactions. Broadly, chemical probing involves reaction of an RNA with a small molecule that is sensitive to the underlying nucleic acid structure^1,2^. Chemical probing has revealed numerous new features of RNA biology and has allowed complex RNA structures to be modeled with good to outstanding accuracy^3–5^. However, a fundamental limitation of most chemical probing strategies is that these experiments merely infer RNA structure from observed reactivity: These approaches do not measure RNA structure directly.

The challenge of directly detecting through-space structural interactions in RNA by chemical probing has been recently addressed by development of the RNA interaction group analyzed by mutational profiling (RING-MaP) technology, in which multiple chemical adducts are visualized in the same RNA strand by a processive relaxed fidelity reverse transcription reaction^6–9^. Normally unreactive or partially reactive nucleotides become transiently exposed to chemical modification through equilibrium fluctuations. Modification of one nucleotide in a dynamically exposed base pair or tertiary interaction blocks local structure refolding, increasing the probability of subsequent modification of the pairing partner (Figure 1A). Through-space structural communication is detected from interdependent modification events in the same molecular strand of RNA. The key advantage of the RING-MaP chemical probing method is thus the ability to directly detect, rather than merely model or predict, RNA duplexes^7,8^, tertiary interactions, and long-range structural communication^6,9,10^ (Figure 1B). To date, RING-MaP correlated chemical probing has been carried out using dimethyl sulfate (DMS).

**Figure 1.**
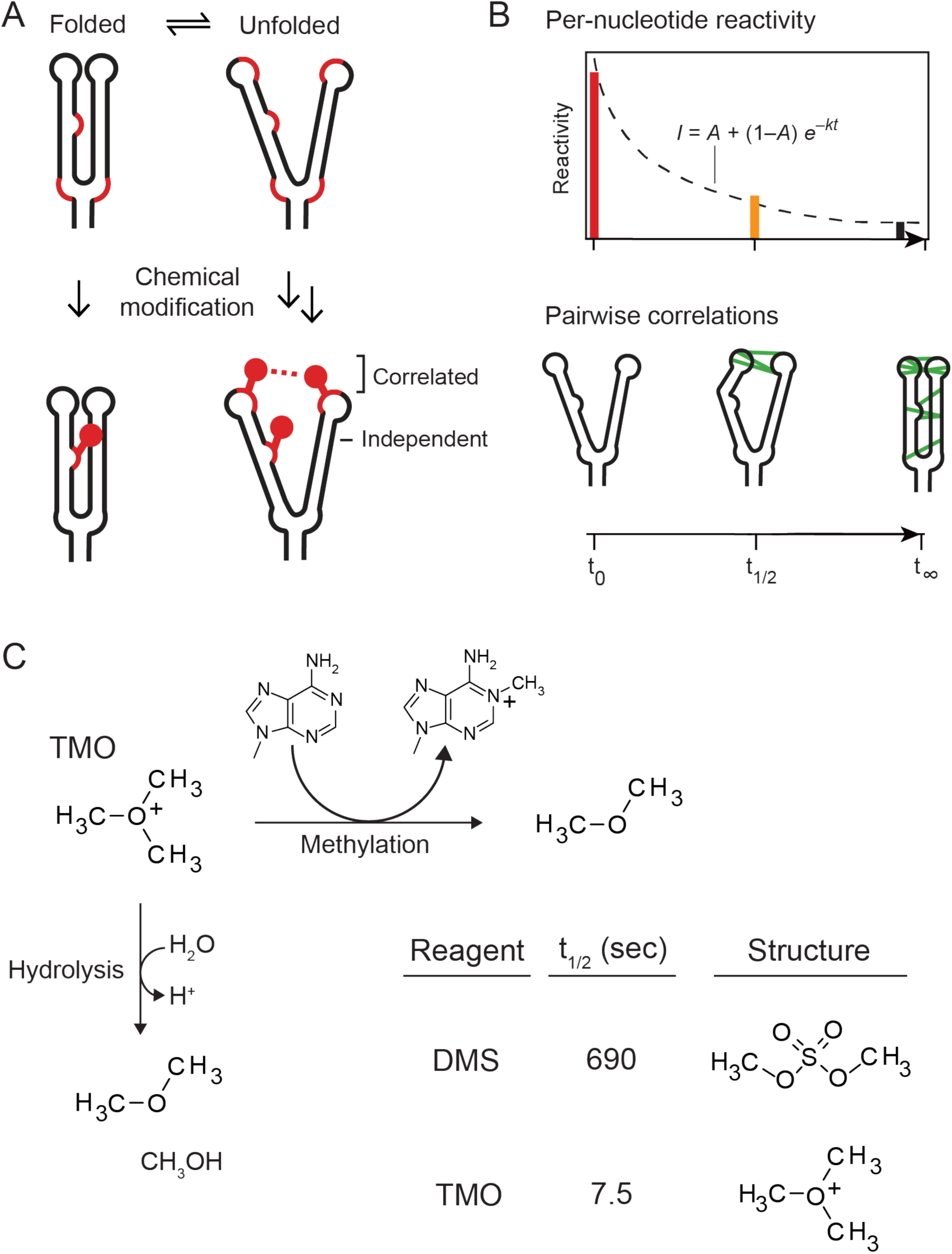
Correlated probing and TMO reactivity. (A) Mechanism of correlated chemical probing. Fluctuations in base-pairing or higher-order structure expose lowly reactive regions to chemical modification, such that one modification event promotes reaction with a second site (correlated adducts). In contrast, modifications to low-structure sites are modified independently of other modification events. Reactive sites are emphasized with red backbone. (B) Strategies for analyzing time-resolved probing data. (top) Time series of per-nucleotide reactivity changes enable kinetic modeling and (bottom) pairwise correlations directly detect through-space RNA interactions. (C) RNA methylation and hydrolysis reaction mechanisms for TMO. Inset shows half-life and structure relative to DMS.

DMS has been used extensively to examine interrelationships between RNA structure and cellular function^1,11^. DMS is valued for its high solubility and its ability to penetrate cell membranes, which enables *in vivo* structure probing^11^. DMS has high reactivity with unpaired adenine and cysteine residues, and recent advances have made it possible to use DMS as a probe for all four ribonucleotides^8^. DMS reacts slowly with RNA, with a reaction half-life of ∼11 min (Figure 1C). DMS probing is therefore limited to interrogating RNA structures at equilibrium and measuring structural cooperativity, but cannot be used to investigate time-resolved processes such as RNA folding.

## Results

We sought to identify a chemical probe with the advantageous features of DMS but with orders of magnitude faster reactivity. After screening a number of reagents, we identified trimethyloxonium tetrafluoroborate (TMO)^12^, which reacts through competing RNA alkylation and self-quenching hydrolysis pathways (Figure 1C). TMO is a small highly soluble reagent, like DMS, but notably differs in that it is a cation. The half-life for the hydrolysis of TMO in aqueous buffered solution, determined by monitoring the time-dependent change in pH upon addition of alkylating agent, is 7.5 sec. Thus, TMO reacts 90 times more rapidly than DMS (Figure 1C, Supporting Figure 1). TMO is therefore appropriate for probing RNA folding processes that occur on second to minute time scales.

We first compared the reactivities of TMO and DMS by probing the structure of the *B. stearothermophilus* ribonuclease P (RNase P) catalytic domain at equilibrium. RNase P has a complex secondary and tertiary structure and its high-resolution structure is known^13,14^. Sites of per-nucleotide chemical modification were identified using the MaP strategy^6,8^. Both TMO and DMS react preferentially with unpaired nucleotides, with high reactivities observed at loops and bulges (Figure 2A and Supporting Figure 2). We observed modest reactivity differences at a subset of nucleotides, likely reflective of differences in probe structure and charge (Figure 2A). Overall, TMO and DMS reactivities are strongly correlated, indicative of similar reactivity preferences for unpaired nucleotides (Figure 2B).

**Figure 2.**
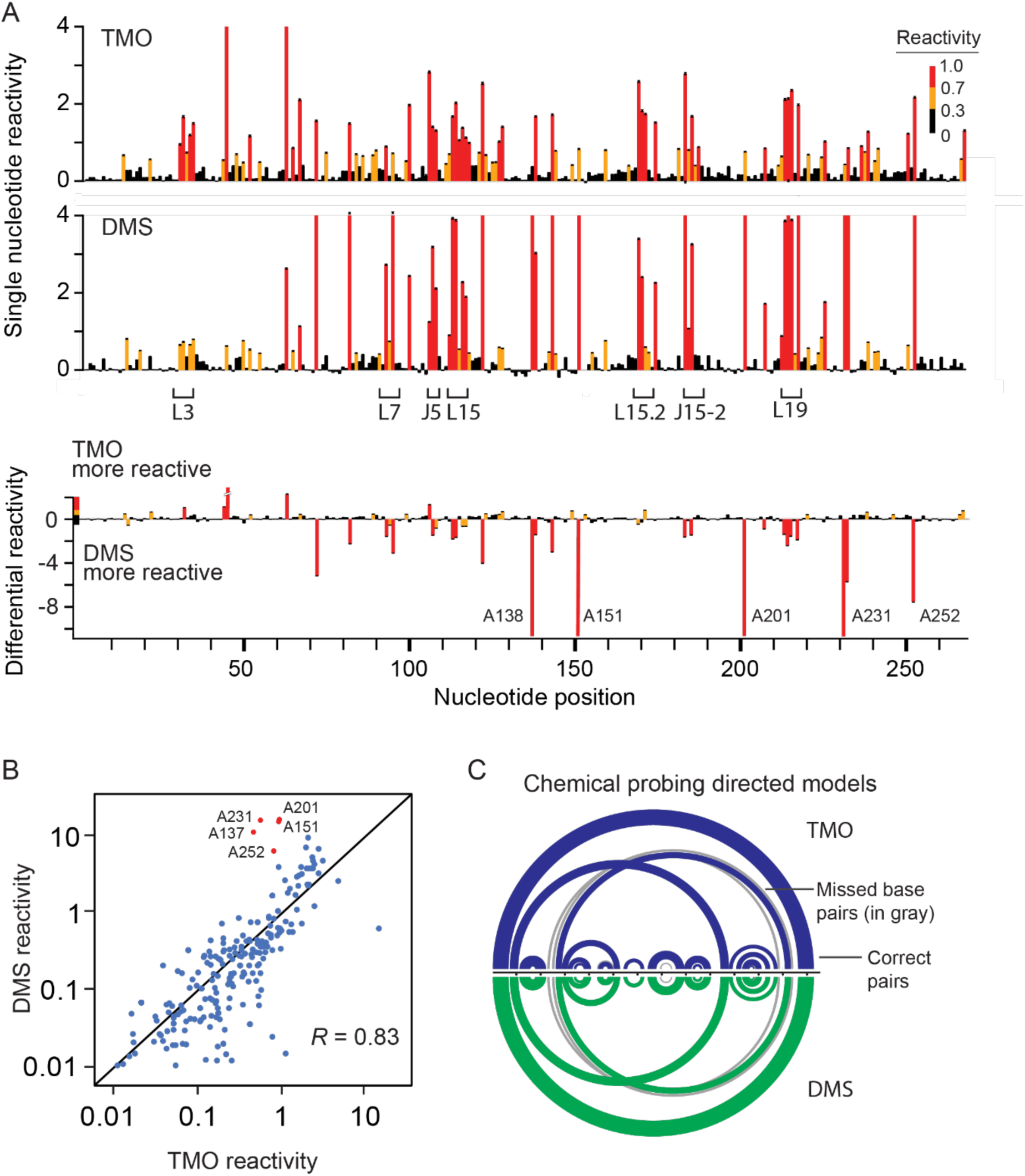
Comparison of TMO and DMS reactivities. (A) Normalized reactivities (top) observed for TMO and DMS for the RNase P catalytic domain. Differential reactivity (bottom) occurs preferentially at sites of noncanonical base pairing (labeled nucleotides). Single-stranded loop (L) and junction (J) landmarks are emphasized. (B) Reactivity correlation for TMO versus DMS; each point indicates an individual nucleotide. Sites that show strong DMS reactivity enhancement are colored red and correspond to non-canonically base paired nucleotides in the accepted secondary structure. (C) Chemical probing-directed secondary structure models for the RNase P catalytic domain based on TMO and DMS reactivities. Structures were modeled using *ShapeKnots*^8^. Arcs indicate base pairs. Blue and green arcs represent correctly modeled base pairs for TMO and DMS, respectively; gray arcs indicate the small number of missed base pairs relative to the accepted structure.

Reactivity profiles can be converted to pseudo-free energy restraints and used to direct RNA secondary structure modeling^8^. DMS reactivity-informed structural modeling of the RNase P accurately recovers both the long- and short-range helices, and the pseudoknot (Figure 2C). TMO reactivity data produce a secondary structure model for RNase P of equivalent accuracy (Figure 2C). Given this ability to model secondary structure with high accuracy and that the TMO reaction has a half-life of 7.5 seconds, we reasoned that TMO could be used to follow time-dependent RNA folding processes at single-nucleotide resolution in two distinct ways: Time-series per-nucleotide reactivity data can be used to observe folding kinetics and, using pair-wise correlation maps, through-space RNA interaction networks can be visualized at each time point (Figure 1B).

We used time-resolved probing with TMO to characterize the folding landscape of an *in vitro* transcript of RNase P. RNA refolding experiments were initiated by addition of Mg^2+^ (to 10 mM) to a buffered solution of RNase P (5 µM, pH 8) preincubated at 37 °C. Aliquots were removed at intervals ranging from 5 to 1200 seconds after Mg^2+^ addition, and were added to one-tenth volume of TMO (to 80 mM final concentration). TMO reactivity profiles were used to generate data-directed structural models of the RNase P RNA as a function of time.

RNase P architecture is composed of canonical base pairing in helices P1-P19, noncanonical pairing and stacking throughout the catalytic core, and a long-range tertiary pairing between loops L5.1 and L15.1 (Figure 3A)^13^. Prior to the addition of Mg^2+^, helices P2, P5, and P15 were reactive and thus not formed. P2, in particular, forms a pseudoknot critical to the tertiary packing of the RNA. In addition, nucleotides in the catalytic core and in the L5.1-L15.1 loop-loop interaction were reactive and thus not formed. (Supporting Figure 2). Time-resolved TMO probing revealed that nucleotides in the P2, P5 and P15 helices refolded slowly upon Mg^2^ addition (Figure 3B). Nucleotides in the catalytic core showed slower reactivity changes than did nucleotides in the P2, P5, and P15 helices. Most intriguingly, loops L5.1 and L15.1 were rapidly protected from TMO reaction after Mg^2^ addition, suggestive of a critical role for this element in the folding landscape. Global structural transitions of the RNase P catalytic domain have been monitored previously by their circular dichroism profile. These prior studies revealed that the RNA folds through a, currently uncharacterized, meta-stable intermediate to form the equilibrium structure^15^. Time-resolved TMO reactivity data suggest that P2, P5, the catalytic core, and the L5.1-L15.1 interaction might all contribute to formation of intermediates in the RNase P RNA folding pathway.

**Figure 3.**
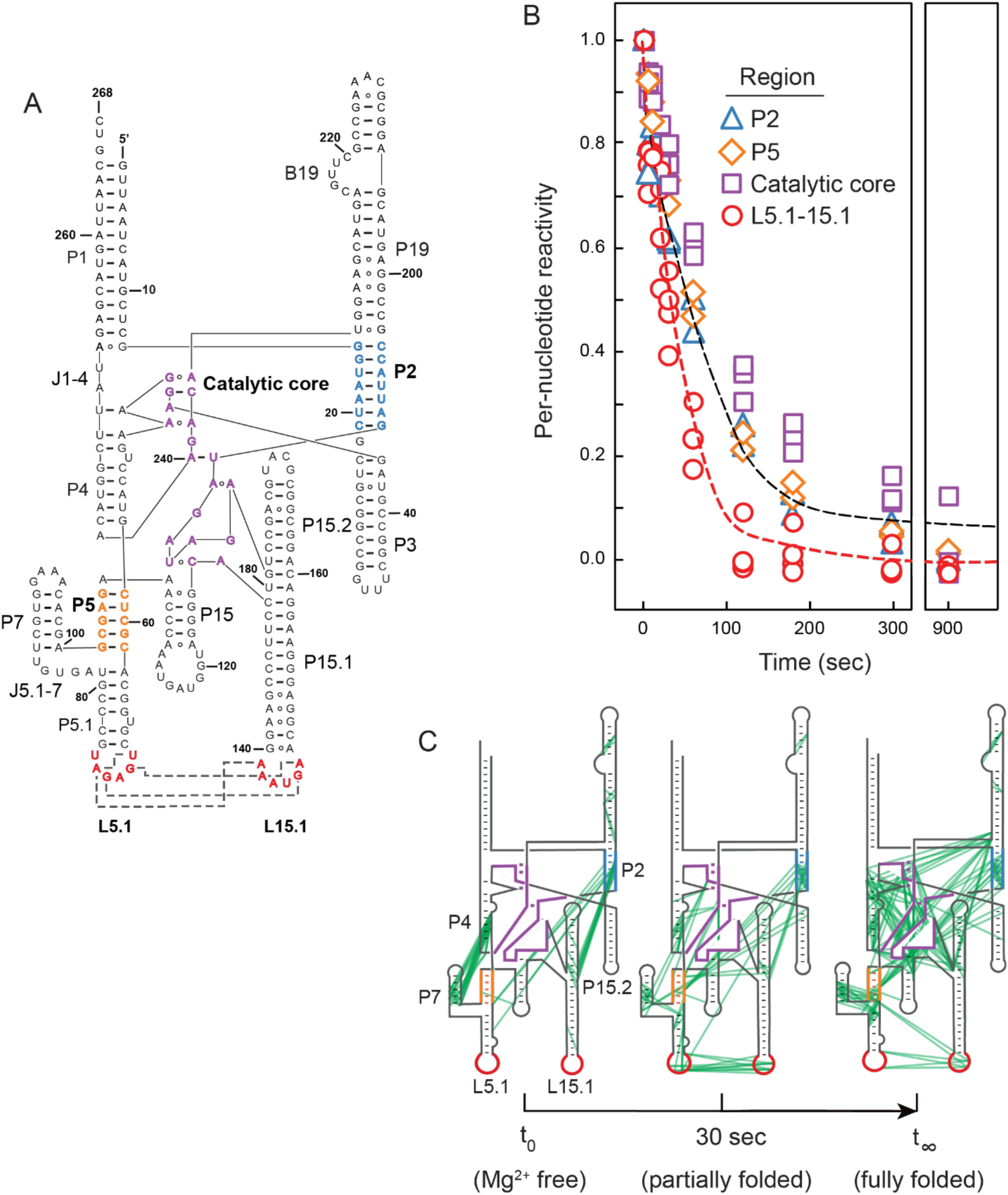
Time-resolved folding of the RNase P catalytic domain. (A) Secondary structure of the RNase P RNA. Regions observed to fold slowly upon Mg^2+^ addition are colored. (B) Time-dependent reactivity profiles for slowly folding domains. Individual per-nucleotide reactivities within domains are shown as points. The red and black lines show best fits to averaged reactivities for nucleotides that form the L5.1-L15.1 interaction and for all other slow folding domains, respectively. (C) Pairwise through-space correlation network for the RNase P folding landscape. Correlations (green lines) are superimposed on the RNase P secondary structure. Slowly folding domains are emphasized with the same color scheme used in panel A.

We next directly visualized through-space structural changes in the RNase P RNA based on pairwise correlations (Figure 1B). Superposition of time-series correlation data onto the secondary structure revealed extensive changes in structural communication as the RNase P folds after Mg^2+^ addition (Figure 3C, green lines). Cross-helix communication is relatively sparse in the absence of Mg^2+^, consistent with the role of Mg^2+^ in stabilizing tertiary RNA structures. In addition, correlations linking P4-P7 and P2-P15.2 were observed in the absence of Mg^2+^ but disappeared in the final structure, indicative of misfolding in the Mg^2+^-free structure. Correlations between the L5.1 and L15.1 loops appeared rapidly upon addition of Mg^2+^, and then decreased gradually as the RNA became fully folded (Figure 3C). The observed decrease in correlation density in L5.1-L15.1 after 60 seconds likely reflects the high stability of this interaction; although this interaction remains detectable, fewer pairwise modifications are observed as these non-canonical interactions form with high stability. In sum, direct visualization of through-space folding (Figure 3C) reveals rapid formation of the tertiary L5.1-L15.1 loop-loop interaction followed by slower folding of the P2 pseudoknot and catalytic core.

Our study thus suggested that the L5.1-L15.1 tertiary interaction plays a critical role in guiding native RNA folding. We therefore examined the role of the L5.1-L15.1 interaction by mutating two nucleotides in L15.1. The mutations extended helix P15.1 by two base pairs and partially disrupted non-canonical tertiary base pairing between L5.1 and L15.1 in the native sequence (Figure 4A). TMO reactivity showed that neither the L5.1-L15.1 tertiary interaction nor the P2-P4 pseudoknot formed fully in the mutant, as these nucleotides were more reactive than for the native sequence RNA (Figure 4B). These reactivity data reveal that the L5.1-L15.1 interaction facilitates pseudoknot formation. RING correlation time series data directly detect, rather than merely infer, the absence of RNA interactions between L5.1 and L15.1 and the absence of the P2 pseudoknot in the mutant RNase P (Figure 4C, Supporting Figure 3). The global effect of disrupting the L5.1-L15.1 loop-loop interaction reveals that this tertiary element plays a critical role in coordinating RNA folding at both base-pairing and tertiary-structure levels.

**Figure 4.**
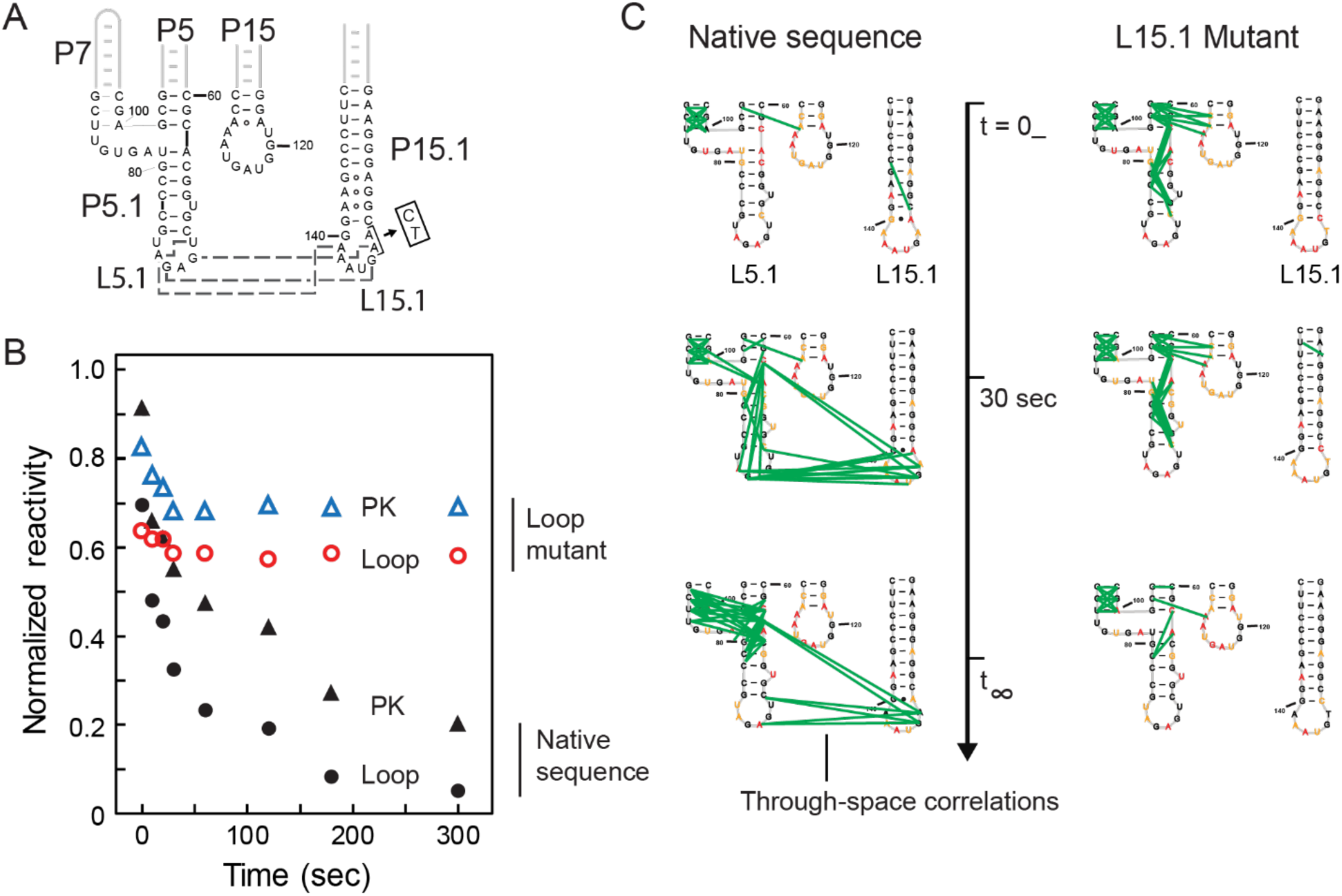
Analysis of RNase P catalytic domain L5.1-L15.1 tertiary interaction on folding. (A) Structure of mutant designed to disrupt the L5.1-L15.1 interaction. Mutations made in L15.1 are indicated by arrow and box. Tertiary contacts (disrupted by the mutant) are shown with gray dashed lines. (B) Reactivities in the pseudoknot (PK) and L5.1 and L15.1 loops as a function of time for the native-sequence and L15.1 mutant RNAs. (C) Time-dependent, correlated chemical probing differences between the native-sequence and mutant RNAs. Green lines represent individual, directly detected, nucleotide-nucleotide interactions.

## Discussion

RNA folding is broadly modeled as hierarchical, meaning that base-paired secondary structures form independently of higher-order tertiary structures, with tertiary interactions occurring after secondary structure formation^16^. Within this framework, structural coupling is also observed such that, when examined at equilibrium, disruption of a tertiary structure interaction also destabilizes secondary structure^17,18^. Using single-molecule correlated probing to visualize the time evolution of dozens of through-space interactions simultaneously (Fig. 3C, 4C), we observed two distinctive features in the RNase P folding reaction. First, folding is clearly not hierarchical: many interactions occurred at early folding time points but then disappear in the final structure, consistent with substantial reorganization of both secondary and tertiary structure. Second, early formation of the long-range L5.1-L15.1 tertiary interaction drives native folding at both the secondary and tertiary structure levels. These complex and non-hierarchical folding patterns were directly visualized as single-molecule pairwise RING correlations.

Time-resolved, correlated chemical probing experiments provide a powerful approach to characterize RNA folding landscapes. TMO is a unique alkylating agent with rapid, but readily controlled, reaction kinetics and high reactivity rates, allowing for layering of single-molecule pairwise correlation data with per-nucleotide reactivity information to directly reveal RNA folding intermediates and pathways. This study moves chemical probing beyond merely inferring dynamics of RNA structure formation, to enable simultaneous and direct measurement of multiple complex folding features for functionally important RNAs.

## Methods

### TMO handling

TMO is a nonvolatile, hygroscopic salt. TMO stocks were handled in a fume hood and stored in desiccator at -20 °C. Due to the high reactivity of TMO, stock solutions of TMO were prepared in 1:2 v/v nitromethane/sulfolane (NS), as follows: TMO was first fully dissolved in nitromethane. Subsequently the TMO solution was diluted with two volumes sulfolane. Final concentrations of TMO stock solutions used in this work (before dilution with RNA) were 1.0 M. Sulfolane is a solid at room temperature; it was warmed to 37 °C immediately prior to use. Due to the fast TMO reaction kinetics, the large volume of RNA solution was added to the smaller TMO volume and mixed by immediate rapid pipetting. Unlike DMS probing, no quench step is needed to stop the TMO reaction.

### Hydrolysis rate determination

Hydrolysis was monitored by adding 200 µL of chemical probe mixture (1.0 M TMO or 1.7 M DMS in NS) to 1800 µL of reaction buffer (11.1 mM MgCl_2_, 111 mM NaCl, 333 mM bicine, pH 8.0) equilibrated at 37 °C. Mixing the components yielded a final concentration of 100 mM TMO or 170 mM DMS with 10 mM MgCl_2_, 100 mM NaCl, 300 mM bicine, pH 8.0, 3.3 v/v% nitromethane, and 6.6 v/v% sulfolane. Pseudo-first order rate constants were obtained by monitoring the pH change over time (PASPORT pH sensor connected to a Pasco interface, sampling frequency of 2 Hz). pH values were normalized to values between 0 and 1, where *pH*_x_ is the *pH* at time x, *pH*_t=∞_ is the final *pH* value, and *pH*_t=0_ is the initial *pH*: 

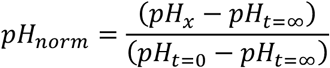

After normalization, hydrolysis rates were determined by fitting a time series to a first-order rate equation.

### RNA synthesis

DNA templates (IDT) encoding the *B. stearothermophilus* RNase P catalytic domain^13^, inserted between 5’ and 3’ structure cassette flanking sequences^19^ and preceded by a 5’ T7 promoter sequence, were amplified by PCR. The DNA template was recovered (Omega Mag-Bind beads) and eluted in water. RNAs were transcribed *in vitro* (2.5 mM each NTP, 25 mM MgCl_2,_ 40 mM Tris, pH 8.0, 2.5 mM spermidine, 0.01% wt/vol Triton X-100, 10 mM DTT, 0.025 units pyrophosphatase, 2.5 µg T7 polymerase, 1.125 µg DNA template; in 50 µL H_2_O; 37 °C; 6 h). RNA was recovered (Omega Mag-Bind beads) and resuspended in 50 µL of 0.1× TE (1 mM Tris, pH 8.0, 0.1 mM EDTA).

### Equilibrium RNA structure probing

RNA (5 pmol) in 6 µL 0.1× TE was heated at 95 °C for 2 min, cooled on ice, and mixed with 3 µL of 3× folding buffer (33 mM MgCl_2_, 333 mM NaCl, 1 M bicine, pH 8). The resulting solution, containing 5 pmol of RNA in 110 mM NaCl, 333 mM bicine, and 11 mM MgCl_2_, was incubated at 37 °C for 20 min. The RNA solution was added to 1 µL of reagent (1.0 M TMO or 1.7 M DMS in NS). The TMO reaction is self-quenching; DMS was quenched with 1 µL of 2-mercaptoethanol after 6 minutes of incubation at 37 °C. The no-reagent control reagent contained 1 µL of NS. Modified RNA was recovered (Omega Mag-Bind beads) and resuspended in 20 µL of 0.1× TE.

### Time-resolved RNA structure probing

RNA (65 pmol) in 52 µL 0.1× TE was heated at 95 °C for 2 min, cooled on ice, and mixed with 39 µL of 3× folding buffer (333 mM NaCl, 1 M bicine, pH 8). The resulting solution, containing 65 pmol of RNA in 110 mM NaCl and 333 mM bicine, was incubated at 37 °C for 10 min. A Mg^2+^-free sample was removed for addition to TMO reagent (100 mM TMO in NS). Tertiary structure folding was initiated by adding 12 µL of 100 mM MgCl_2_ (final Mg^2+^ concentration was 10 mM). After mixing, 9 µL of RNA solution was removed (at 5, 10, 15, 20, 30, 60, 120, 180, 300, 600, 900, and 1200 s) and added directly into 1 µL of 10× reagent (1.0 M TMO in 1:2 v/v NS). For the no-reagent control, the aliquot was added to 1 µL 1:2 v/v NS. Modified RNA was recovered with Omega Mag-Bind beads and resuspended in 20 µL of 0.1× TE (1 mM Tris pH 8.0, 0.1 mM EDTA).

### Reverse transcription

MaP was performed essentially as described^7,20^. In brief, a 10 µL solution containing RNA, 200 nM gene specific primer (Table S1), and 2 mM premixed dNTPs was incubated at 65 °C for 5 min followed by incubation at 4 °C for 2 min. To the solution was added 9 µL of 2.22× MaP buffer (1× MaP buffer contains 1M betaine, 50 mM Tris, pH 8.0, 75 mM KCl, 10 mM DTT, 6 mM MnCl_2_), and the mixture was incubated at room temperature for 2 min. SuperScript II Reverse Transcriptase (1 µL, Invitrogen) was added, and reverse transcription was performed according to the following temperature program: 25 °C for 10 min, 42 °C for 90 min, 10 × [50 °C for 2 min, 42 °C for 2 min], 72 °C for 10 min. cDNA was then purified (Illustra MicroSpin G-50 columns, GE Healthcare).

### Library preparation and sequencing

Sequencing libraries were prepared from cDNA products using a two-step PCR approach (Table S1)^8^. In the first PCR step, a 5-µL aliquot of purified cDNA was amplified for Illumina sequencing with the following temperature program: 98 °C for 30 s, 15 × [98 °C for 5 s, 68 °C for 20 s, 72 °C for 20 s], 72 °C for 2 min. The PCR product was recovered (Omega Mag-Bind beads) and eluted in water. In the second PCR step, treatment-specific barcodes were added to the ends of amplicons, with the following temperature program: 98 °C for 30 s, 15 × [98 °C for 5 s, 68 °C for 20s, 72 °C for 20s], 72 °C for 2 min. PCR products were recovered (Omega Mag-Bind beads), pooled, and sequenced (Illumina MiSeq instrument; 500 cycle kit).

### Sequence alignment and mutation parsing

*ShapeMapper* (v2.1.4) was used to align reads to the reference sequence and to identify positions mutated during MaP^21^. The --output-parsed option was used to generate mutational files that serve as input to *PairMapper* and *RingMapper* correlation analysis software packages^8^.

### Rate determinations

Per-nucleotide reactivities were processed using a custom Python package. First, a time-series matrix of per-nucleotide reactivities was created from individual reactivity profiles output by *ShapeMapper*. Reactivities were normalized for each nucleotide position by computing reactivity ratios relative to the initial reactivity *R*_1_ and the final reactivity *R*_2_ across the time series: 

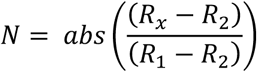

Normalized nucleotide reactivities were input to the *scipy* curve fitting algorithm as a function of time. Nucleotide reactivities were fit to a single exponential: 

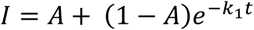

where *I* is normalized reactivity, *A* is a reactivity amplitude, and *k*_1_ is the rate constant. Nucleotides with curve fit correlations greater than 0.9 were analyzed further. Fitted kinetic curve plots were visually inspected for goodness of fit (JMP Pro 14).

### Correlation analysis

*RingMapper* and *PairMapper* were used to compute correlated modification events^8^. *PairMapper* and *RingMapper* were run with default settings. *RingMapper* outputs a table containing correlation position, read depth, and correlation significance. Negative correlations were removed. Those with positive correlations were required to have average product corrected (APC) G statistic strength above 100; if this criterion was not met, the events were removed. The average product corrected G statistic is a metric of correlation significance normalized relative to background correlation significance. *PairMapper* and *RingMapper* software packages are available for download at https://github.com/Weeks-UNC.

### RNase P catalytic domain structure modeling

*ShapeKnots* (from software package *RNAstructure*, v6.1)^22^ was used to model RNase P secondary structure. Required parameters were the RNase P catalytic domain primary sequence^13^ and the output .ct file name. Pseudo-free energy restraints were generated by calling the flag *-dmsnt* to use TMO- or DMS-normalized reactivities and the flag *-x* was called to incorporate *PairMapper* .bp files, thus introducing two TMO- or DMS-based bonuses (per-nucleotide and pairwise correlation) into the secondary structure calculation^8^. The *-m* 1 option was used to select only the minimum free energy structure, outputting a .ct file containing secondary structure information.

## Supporting Information

One table with RNA sequence information, and three figures.

## Acknowledgements

We thank A.M. Mustoe for insightful discussions and for ongoing development of the *RingMapper* and *PairMapper* software.

## References

(1) Peattie, D. A.; Gilbert, W. Chemical probes for higher order structure in RNA. Proc. Natl. Acad. Sci. USA 1980, 77, 4679–4682.

(2) Weeks, K. M. Advances in RNA structure analysis by chemical probing. Curr. Opin. Struct. Biol. 2010, 20, 295–304.

(3) Smola, M. J.; Rice, G. M.; Busan, S.; Siegfried, N. A.; Weeks, K. M. Selective 2’-hydroxyl acylation analyzed by primer extension and mutational profiling (SHAPE-MaP) for direct, versatile and accurate RNA structure analysis. Nat. Protoc. 2015, 10, 1643–1669.

(4) Mustoe, A. M.; Corley, M.; Laederach, A.; Weeks, K. M. Messenger RNA Structure Regulates Translation Initiation: A Mechanism Exploited from Bacteria to Humans. Biochemistry 2018, 57, 3537–3539.

(5) Boerneke, M. A.; Ehrhardt, J. E.; Weeks, K. M. Physical and Functional Analysis of Viral RNA Genomes by SHAPE. Annu. Rev. Virol. 2019, 6, 93–117.

(6) Homan, P. J.; Favorov, O. V; Lavender, C. A.; Kursun, O.; Ge, X.; Busan, S.; Dokholyan, N. V; Weeks, K. M. Single-molecule correlated chemical probing of RNA. Proc. Natl. Acad. Sci. U. S. A. 2014, 111, 13858–13863.

(7) Krokhotin, A.; Mustoe, A. M.; Weeks, K. M.; Dokholyan, N. V. Direct identification of base-paired RNA nucleotides by correlated chemical probing. RNA 2017, 23, 6–13.

(8) Mustoe, A. M.; Lama, N. N.; Irving, P. S.; Olson, S. W.; Weeks, K. M. RNA base-pairing complexity in living cells visualized by correlated chemical probing. Proc. Natl. Acad. Sci. U. S. A. 2019, 116, 24574–24582.

(9) Sengupta, A.; Rice, G. M.; Weeks, K. M. Single-molecule correlated chemical probing reveals large-scale structural communication in the ribosome and the mechanism of the antibiotic spectinomycin in living cells. PLOS Biol. 2019, 17, e3000393.

(10) Dethoff, E. A.; Boerneke, M. A.; Gokhale, N. S.; Muhire, B. M.; Martin, D. P.; Sacco, M. T.; McFadden, M. J.; Weinstein, J. B.; Messer, W. B.; Horner, S. M.; Weeks, K. M. Pervasive tertiary structure in the dengue virus RNA genome. Proc. Natl. Acad. Sci. U. S. A. 2018, 115, 11513–11518.

(11) Wells, S. E.; Hughes, J. M. X.; Ares, M. Use of dimethylsulfate to probe RNA structure in vivo. Methods Enzymol. 2000, 318, 479–493.

(12) Stahl, I.; Seapy, D. G. Trimethyloxonium Tetrafluoroborate. In Encyclopedia of Reagents for Organic Synthesis; John Wiley & Sons, Ltd: Chichester, UK, 2008.

(13) Kazantsev, A. V; Krivenko, A. A.; Pace, N. R. Mapping metal-binding sites in the catalytic domain of bacterial RNase P RNA. RNA 2009, 15, 266–276.

(14) Kazantsev, A. V.; Rambo, R. P.; Karimpour, S.; Santalucia, J.; Tainer, J. A.; Pace, N. R. Solution structure of RNase P RNA. RNA 2011, 17, 1159–1171.

(15) Fang, X. W.; Golden, B. L.; Littrell, K.; Shelton, V.; Thiyagarajan, P.; Pan, T.; Sosnick, T. R. The thermodynamic origin of the stability of a thermophilic ribozyme. Proc. Natl. Acad. Sci. U. S. A. 2001, 98, 4355–4360.

(16) Brion, P.; Westhof, E. Hierarchy and dynamics of RNA folding. Annu. Rev. Biophys. Biomol. Struct. 1997, 26, 113–137.

(17) Woodson, S. A. Compact Intermediates in RNA Folding. Annu. Rev. Biophys. 2010, 39, 61–77.

(18) Herschlag, D.; Bonilla, S.; Bisaria, N. The story of RNA folding, as told in epochs. Cold Spring Harb. Perspect. Biol. 2018, 10, a032433.

(19) Merino, E. J.; Wilkinson, K. A.; Coughlan, J. L.; Weeks, K. M. RNA structure analysis at single nucleotide resolution by Selective 2′-Hydroxyl Acylation and Primer Extension (SHAPE). J. Am. Chem. Soc. 2005, 127, 4223–4231.

(20) Siegfried, N. A.; Busan, S.; Rice, G. M.; Nelson, J. A. E.; Weeks, K. M. RNA motif discovery by SHAPE and mutational profiling (SHAPE-MaP). Nat. Methods 2014, 11, 959–965.

(21) Busan, S.; Weeks, K. M. Accurate detection of chemical modifications in RNA by mutational profiling (MaP) with ShapeMapper 2. RNA 2018, 24, 143–148.

(22) Reuter, J. S.; Mathews, D. H. RNAstructure: software for RNA secondary structure prediction and analysis. BMC Bioinformatics 2010, 11, 129.

