## Supplemental Table and Figures for "Time-resolved, single-molecule, correlated chemical probing of RNA"

Jeffrey E. Ehrhardt and Kevin M. Weeks
Department of Chemistry
University of North Carolina

**Table S1.** Template sequences and primers used in this study

| RNA | Sequence 5' → 3' |
|---|---|
| RNase P catalytic domain, native sequence | Template:<br>TAATACGACTCACTATAGGGCCTTCGGGCCAAGTTAATCATGCTCGGGTAAT<br>CGCTGCGGCCGGTTTCGGCCGTAGAGGAAAGTCCATGCTCGCACGGTGCT<br>GAGATGCCCGTAGTGTTCGTGGAAACACGAGCGAGAAACCCAAATGATGGT<br>AGGGGCACCTTCCCGAAGGAAATGAACGGAGGGAAGGACAGGCGGCGCAT<br>GCAGCCTGTAGATAGATGATTACCGCCGGAGTACGAGGCGCAAAGCCGCTT<br>GCAGTACGAAGGTACAGAACATGGCTTATAGAGCATGATTAACGTCTCGATC<br>CGGTTCGCCGGATCCAAATCGGGCTTCGGTCCGGTTC<br><br>Forward PCR template primer:<br>TAATACGACTCACTATAGGGCCTTCGGG<br><br>Reverse PCR template primer:<br>GAACCGGACCGAAGCCCG<br><br>Reverse Transcription primer:<br>*Same as Step 1 reverse primer<br><br>Step 1 forward primer:<br>CCCTACACGACGCTCTTCCGATCTNNNNNGGCCTTCGGGCCAAGGA<br><br>Step 1 reverse primer:<br>GACTGGAGTTCAGACGTGTGCTCTTCCGATCTNNNNNTTGAACCGGACCGA<br>AGCCCGATTT |
| RNase P, loop mutant | Template:<br>TAATACGACTCACTATAGGGCCTTCGGGCCAAGTTAATCATGCTCGGGTAAT<br>CGCTGCGGCCGGTTTCGGCCGTAGAGGAAAGTCCATGCTCGCACGGTGCT<br>GAGATGCCCGTAGTGTTCGTGGAAACACGAGCGAGAAACCCAAATGATGGT<br>AGGGGCACCTTCCCGAAGGAAATGAACGGAGGGAAGGACAGGCGGCGCAT<br>GCAGCCTGTAGATAGATGATTACCGCCGGAGTACGAGGCGCAAAGCCGCTT<br>GCAGTACGAAGGTACAGAACATGGCTTATAGAGCATGATTAACGTCTCGATC<br>CGGTTCGCCGGATCCAAATCGGGCTTCGGTCCGGTTC<br><br>*Primers for the mutant RNase P are identical to native sequence primers. |

**Figure S1.** Hydrolysis time courses for TMO and DMS at 37 °C. Reagent hydrolysis was monitored by pH.

**Figure S2.** Single-nucleotide TMO reactivity data as a function of position on the secondary structure of the RNase P catalytic domain. TMO reactivities at time points 0 ($Mg^{2+}$ free), 30, 120, and 1200 seconds are superimposed on the equilibrium structure of the RNase P catalytic domain. Red, yellow, and black nucleotides represent high, medium, and low TMO reactivities, respectively (see legend). Dashes connecting nucleotides indicate Watson-Crick base pairing; circles linking nucleotides emphasize non-canonical base pairing.

**Figure S3.** Time-dependent, single molecule, correlated chemical probing of the native-sequence and mutant RNase P RNAs. Correlations (green lines) are superimposed on the RNase P secondary structure. Slowly folding domains are emphasized in color (as per Figure 4). Pseudoknot helices P2 and P5, blue and orange; catalytic core, purple; L5.1-L15.1, red.

**Figure S1**

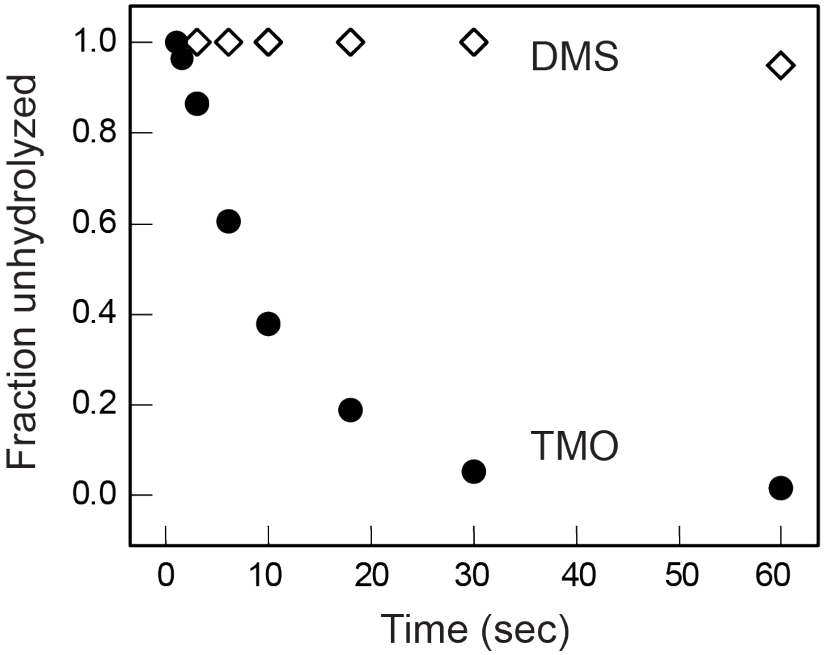

Fraction unhydrolyzed
1.0
0.8
0.6
0.4
0.2
0.0
DMS
TMO
0
10
20
30
40
50
60
Time (sec)

**Figure S2**

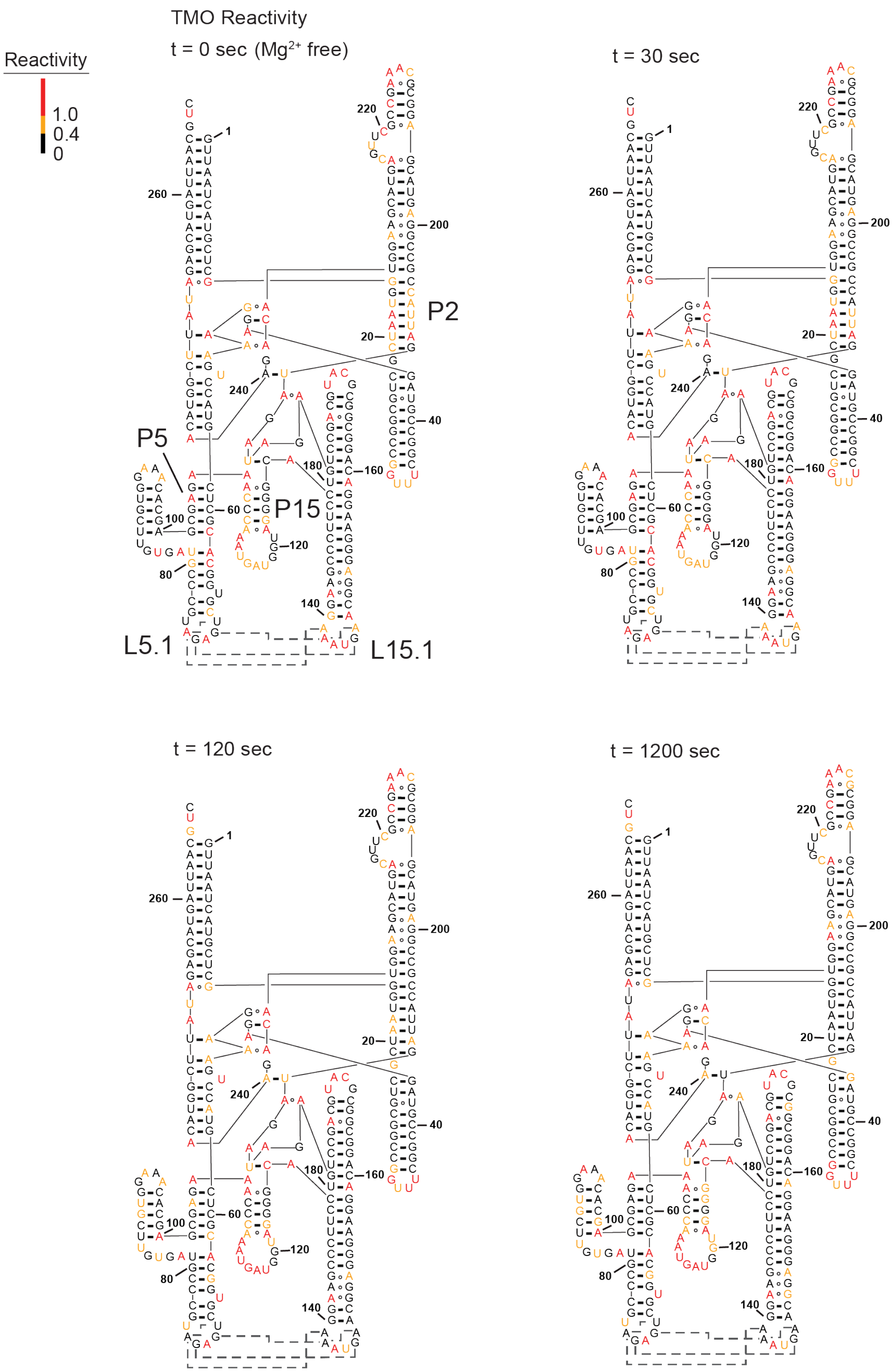
TMO Reactivity
t = 0 sec (Mg2+ free)
Reactivity
1.0
0.4
0
P2
P5
P15
L5.1
L15.1
t = 30 sec
t = 120 sec
t = 1200 sec

**Figure S3**

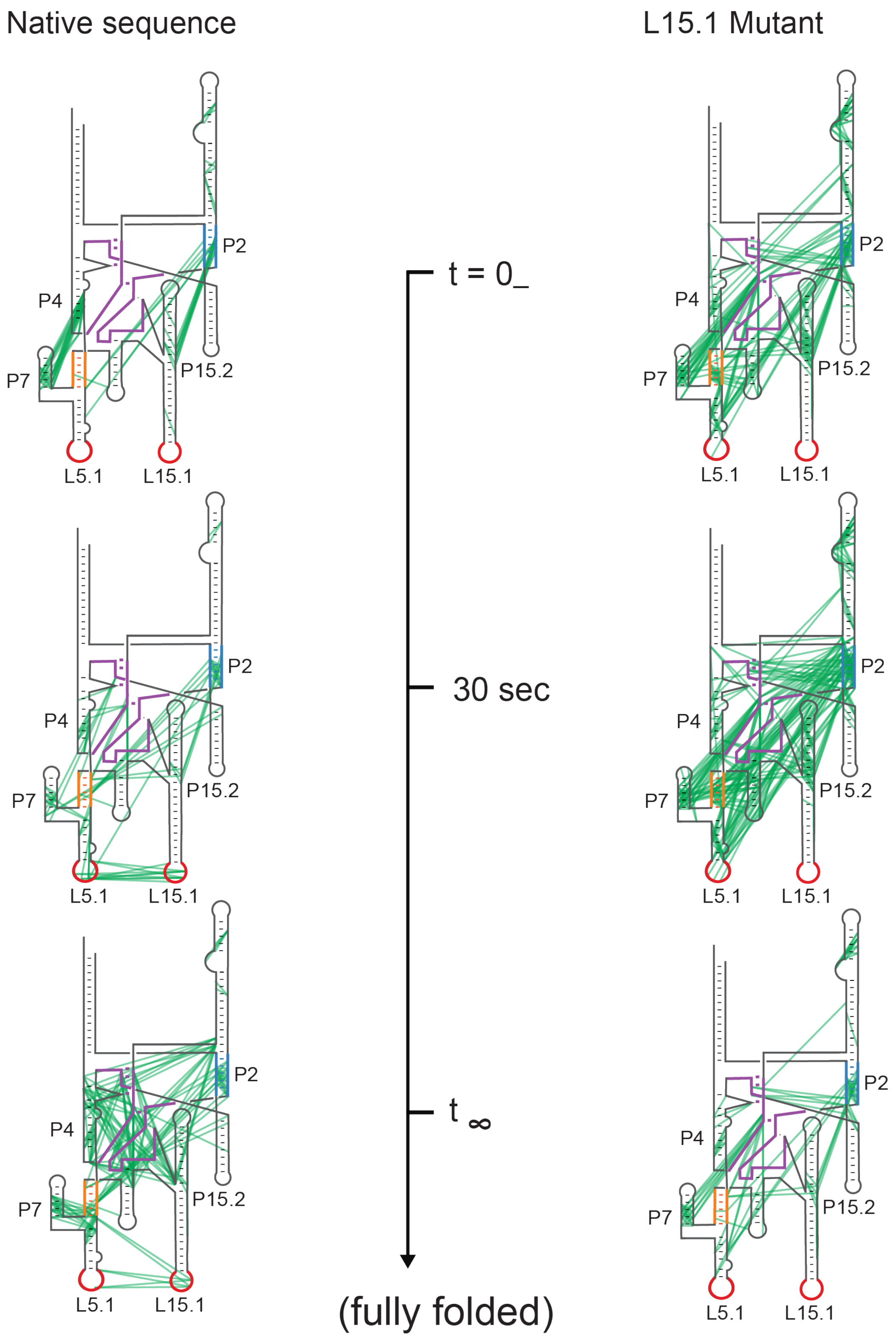
Native sequence
L15.1 Mutant
P2
P4
P7
P15.2
L5.1
L15.1
t = 0_
30 sec
t ∞
(fully folded)
